# Ceramides disrupt hexokinase HKI-VDAC complex assembly and modulate VDAC tilting

**DOI:** 10.64898/2026.04.25.720789

**Authors:** Michael Timme, Joost C. M. Holthuis, Manuel N. Melo

**Affiliations:** Division of Molecular Cell Biology, Department of Biology/Chemistry, Osnabrück University, 49076 Osnabrück, Germany; Center for Cellular Nanoanalytics, Osnabrück University, 49076 Osnabrück, Germany; Instituto de Tecnologia Química e Biológica António Xavier, Universidade Nova de Lisboa, Av. da República, 2780-157 Oeiras, Portugal

## Abstract

Ceramides bind voltage-dependent anion channels (VDACs) to promote mitochondrial apoptosis. However, the underlying mechanism is unclear. Ceramide binding relies on a charged, bilayer-facing glutamate that also controls complex formation between VDACs and pro-survival hexokinase HKI, suggesting that ceramides may exert their pro-apoptotic activity by displacing HKI from VDACs. Here, we challenged this model using coarse-grain molecular dynamics simulations. We show that ceramides impair VDAC-HKI complex assembly, but not via direct competition with HKI for binding to the crucial glutamate. Instead, we find that ceramides disrupt the membrane thinning capacity of polar channel residues adjacent to the glutamate, thereby congesting a low-energy passageway used by HKI for efficient complex formation. We also demonstrate that VDACs are tilted in the membrane and that ceramides act as modulators of channel tilt by neutralizing the hydrophobic mismatch imposed by the polar channel residues. We postulate that this ceramide action further modulates the oligomerization propensity of VDACs. Altogether, our data reveal a mechanistic basis for how ceramides may trigger apoptotic cell death.

## INTRODUCTION

Ceramides are central intermediates of sphingolipid metabolism that are frequently identified as critical mediators of stress-induced mitochondrial apoptosis^1,2^. Numerous studies indicate that cellular ceramides levels rise concomitantly with apoptosis induction in response to stress stimuli, including exposure to TNFα, chemotherapeutic agents, and ionizing radiation^3–6^. Interventions that suppress ceramide accumulation renders cancer cells resistant to these stress stimuli. While these findings support the notion that ceramides are bona fide pro-apoptotic signaling molecules, the mechanisms by which they exert their pro-apoptotic activities are incompletely understood.

Several studies indicate that ceramides can act directly on mitochondria to initiate permeabilization of the outer mitochondrial membrane (OMM) for cytochrome *c*, a point of no return in the apoptotic program that results in an ordered, caspase-mediated self-destruction of cells^7^. For instance, mitochondrial targeting of a bacterial sphingomyelinase to generate ceramides in mitochondria or directing CERT-mediated ceramide transport to mitochondria triggers mitochondrial translocation of Bax, cytochrome *c* release and apoptotic cell death^8–10^. Moreover, mitochondria-specific photorelease of ceramides induces cleavage of Casp9 and PARP1, signifying activation of the intrinsic apoptotic pathway^11^. How a rise in mitochondrial ceramides leads to permeabilization of the OMM to trigger apoptotic cell death remains a subject of debate. Some studies suggest that ceramides in the OMM form channels that are large enough to allow passage of cytochrome *c*^12^. An alternative model postulates that ceramides generated in the OMM of irradiated cells form microdomains that facilitate functionalization of Bax into cytochrome *c*-conducting pores^13^.

Using a chemical screen for ceramide binding proteins combined with coarse-grain molecular dynamics (CG-MD) simulations and functional studies in cancer cells, we previously identified the voltage-dependent anion channels VDAC1 and VDAC2 as core components of a pathway by which ceramides may exert their pro-apoptotic activities^14^. Besides their role as main gatekeepers for the passage of ions and metabolites across the OMM^15^, VDAC channels function as scramblases that mediate phospholipid import into mitochondria^16^. Moreover, VDACs serve as dynamic translocation platforms for Bax and other proteins that control the permeability of the OMM for cytochrome *c* to either initiate or block mitochondrial apoptosis^17–21^. We found that ceramide binding to VDACs critically relies on a uniquely positioned charged glutamate on the outer channel wall that mediates direct contact with the ceramide head group in the bilayer interior^14^. Importantly, substitution of this residue in VDAC2 abolished ceramide binding and rendered human colon cancer cells resistant to ceramide-induced apoptosis. In a subsequent study, we found that the same glutamate is essential for mitochondrial recruitment of hexokinase HKI^22^. Binding of HKI to mitochondrial VDACs has far reaching physiological implications and is thought to promote cell proliferation and survival in rapidly growing hyperglycolytic tumors^20,23,24^.

The notion that HKI and ceramides share a common binding site on VDACs points at a potential mechanism by which ceramides may exert their anti-neoplastic activities. We postulated that ceramides might increase cellular susceptibility to stress-induced apoptosis by impairing the formation of pro-survival HKI-VDAC complexes through direct competition with HKI for binding to the charged, membrane buried glutamate. In this study, we set out to challenge this model using CG-MD simulations.

## RESULTS

### VDAC-HKI complex assembly is perturbed by ceramide

The *N*-terminal α-helix of HKI comprising residues 1-18 (HKI-N) is essential and sufficient for VDAC binding. Using CG simulations, we previously found that VDAC1 and VDAC2 bind HKI-N, provided that the membrane-buried Glu on the channel wall (E73 in VDAC1; E84 in VDAC2) is in a deprotonated, charged state and that the C-termini of the channels are oriented toward the IMS^22^. To investigate whether ceramides affect VDAC-HKI complex formation, we simulated VDAC1 and VDAC2 in an OMM-mimicking bilayer containing different concentrations of C_16:0_ ceramide and monitored binding of HKI-N to the charged, bilayer-facing Glu. In the absence of ceramide, and in line with our previous findings, the *N*-terminal half of HKI-N inserted vertically into the cytosolic leaflet along one side of the channel wall and made intimate contacts with E73^−^ in VDAC1 (**Fig. 1a, b**) and E84^−^ in VDAC2 (data now shown). These binding events were often detected multiple times per simulation and reached µs durations (**Fig. 1a**). The protonated HKI-N-Met1 had the highest contact prevalence with E73^−^ in VDAC1 and E84^−^ in VDAC2 of all HKI-N/VDAC residue pairs (**Fig. 1c-g**). Addition of C_16:0_ ceramide perturbed these interactions in a concentration-dependent manner, with HKI-N binding to both channel isomers almost completely abolished in the presence of 20 mol% C_16:0_ ceramide (Cer; **Fig. 1c-g**). When measuring the contact lifetimes of HKI-N-Met1 with E73^−^ in VDAC1 or E84^−^ in VDAC2 under these conditions, we found that the average contact duration was unaffected (**Suppl. Fig. 1a, b**), indicating that C_16:0_ ceramide primarily interferes with complex assembly rather than promoting complex dissociation. In contrast, HKI-N binding to VDACs was largely unaffected or even enhanced in the presence of 20 mol% C_16:0_ sphingomyelin (SM; **Fig. 1c-g**). C_16:0_ sphingomyelin shares the same backbone as C_16:0_ ceramide but sports a bulkier and more polar headgroup, and is used in this work as a control that ceramide-mediated effects are specific to it and not merely a consequence of increasing the amount of saturated, stiffer tail species. From this, we conclude that ceramide is a specific inhibitor of VDAC-HKI complex assembly.

**Figure 1.**
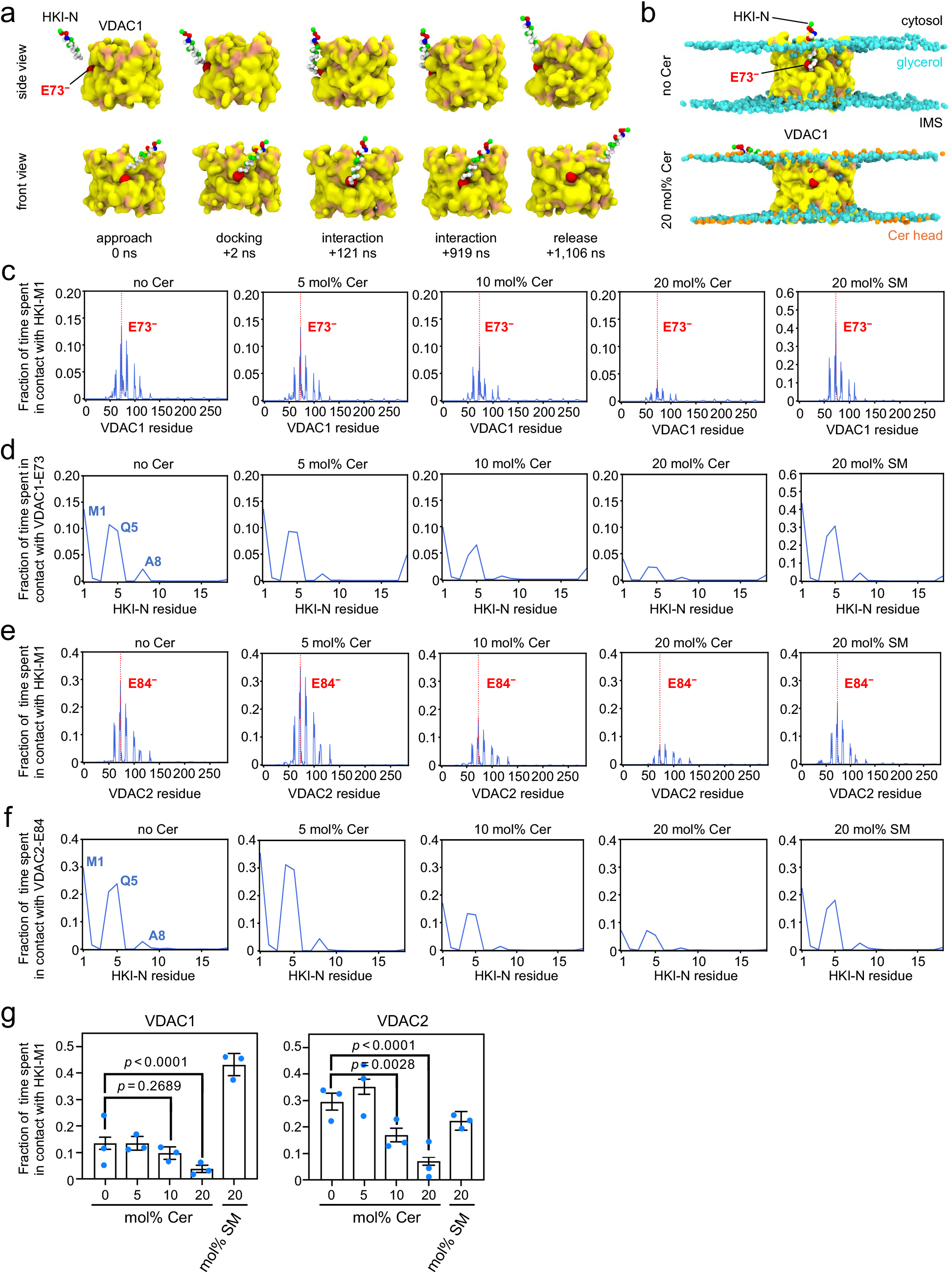
Ceramides disrupt HKI-N binding to VDACs. **(a)** Stills from an MD simulation showing the approach and binding of HKI-N to VDAC1 with a charged (deprotonated) E73 (*red*) and IMS-facing *C*-terminus in an OMM-mimicking bilayer. For clarity, the membrane bilayer is not shown. **(b)** Stills from MD simulations of HKI-N and VDAC1 with a charged E73 (*red*) in an OMM-mimicking bilayer containing either no (*top*) or 20 mol% C_16:0_ ceramide (*bottom*). Phospholipid glycerol backbones and ceramide headgroups are marked in *cyan* and *orange*, respectively. **(c)** Time fraction of contacts between HKI-Met1 and specific residues of VDAC1 with a charged E73. Simulations were performed in an OMM-mimicking bilayer containing the indicated amount of C_16:0_ ceramide (Cer) or C_16:0_ sphingomyelin (SM). Shown are the combined data of three individual replicas with a total simulation time between 217 µs and 279 µs per condition. **(d)** Time fraction of contacts between VDAC1-E73 and specific residues of HKI-N in simulations as in (c). **(e)** Time fraction of contacts between HKI-Met1 and specific residues of VDAC2 in simulations carried out as in (c). Shown are the combined data of three individual replicas with a total simulation time between 217 µs and 277 µs per condition. **(f)** Time fraction of contacts between VDAC2-E84 and specific residues of HKI-N in simulations as in (e). **(g)** Time fraction of contacts between HKI-Met1 and VDAC1-E73. Plotted are the individual replicates from the same simulations as in (c). Data are means ± SEM, n = 3 replicas per condition. SEMs were determined from block averaging over all replica trajectories, with chosen block lengths exceeding the autocorrelation time of bound/unbound states to generate statistically independent samples. *p* values were calculated by unpaired two-tailed *t* test. **(h)** Time fraction of contacts between HKI-Met1 and VDAC2-E84. Plotted are the individual replicates from the same simulations as in (e). Data are means ± SEM, n = 3 per condition, with SEMs and *p* values determined as in (g).

### Ceramide affects HKI binding to VDACs by restricting access to the bilayer-facing Glu

To explore how C_16:0_ ceramide exerts its inhibitory effect on VDAC-HKI complex formation, we next mapped its relative occupancy in MD simulations of VDAC1 in an OMM-mimicking bilayer. This revealed that C_16:0_ ceramide avoids the membrane area surrounding the channel wall, irrespective of its concentration (**Suppl. Fig. 2**). We postulate that a hydrophobic mismatch between the channel and membrane bilayer^25^ (see also below) renders this area energetically unfavorable for C_16:0_ ceramide, a lipid with strong bilayer rigidifying properties^26^. However, C_16:0_ ceramide preferentially populated several discrete membrane regions adjacent to the channel wall, including one directly adjacent to the charged bilayer-facing Glu (E73^−^). This latter region was abolished upon protonation of E73, consistently with our previous findings that a charged E73 is a critical determinant of ceramide–VDAC1 binding^14^. In contrast, C_16:0_ sphingomyelin did not show any enhanced occupancy in this region, regardless of the protonation state of E73 (**Suppl. Fig. 2**).

**Figure 2.**
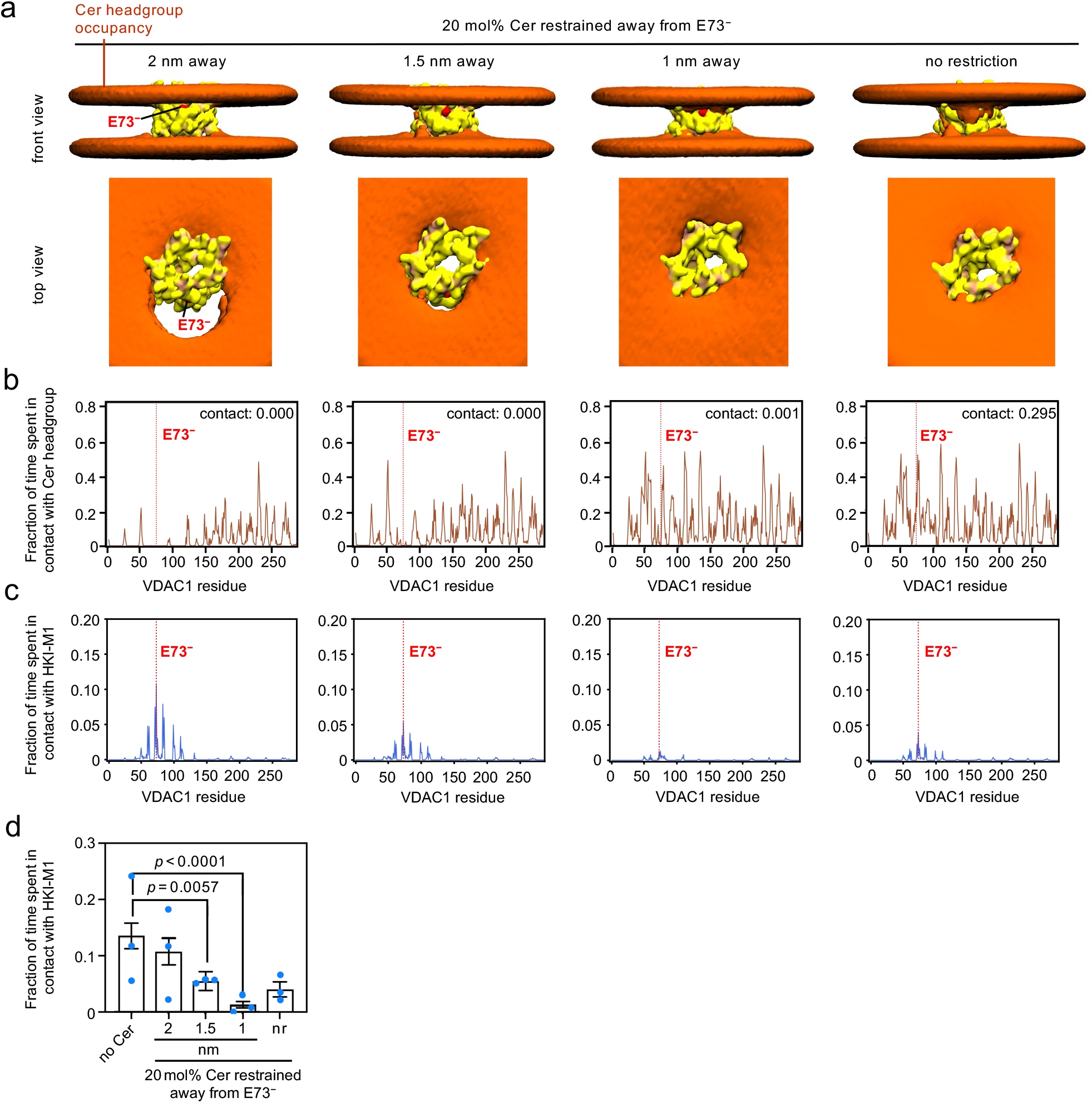
Ceramides modulate HKI-N binding to VDACs by controlling its access to the bilayer-facing Glu. **(a)** Top and front views of MD simulations of VDAC1 with a charged E73 (*red*) and HKI-N in an OMM-mimicking bilayer containing 20 mol% of C_16:0_ ceramide (Cer). A harmonic minimum distance restraint was placed at the indicated distances between each ceramide headgroup and the bilayer-facing E73 side-chain (*red sphere*). Ceramide headgroup occupancies over the course of the simulations are shown as *orange* surfaces with a threshold of 0.05% occupation time or greater. Shown are the combined data of three individual replicas with a total simulation time between 203 µs and 269 µs per condition. For clarity, the occupancies of HKI-N are not shown. **(b)** Time fraction of contacts between ceramide headgroups and specific residues of VDAC1 in simulations as in (a). The relative contact time with E73 per condition is indicated. **(c)** Time fraction of contacts between HKI-Met1 and specific residues of VDAC1 in simulations as in (a). **(d)** Time fraction of contacts between HKI-Met1 and VDAC1-E73. Plotted are the individual replicates from the same simulations as in (a) with replicates from MD simulations carried out in the absence of ceramides serving as a reference. Data are means ± SEM, n = 3 per condition, with SEMs and *p* values determined as in **Fig. 1g**.

The foregoing raised the possibility that C_16:0_ ceramide disrupts VDAC-HKI complex formation by directly competing with HKI-N for binding to the charged, bilayer-facing Glu. To challenge this idea, we simulated VDAC1 with HKI-N in an OMM-mimicking bilayer containing 20 mol% C_16:0_ ceramide and placed a harmonic minimum distance restraint ranging from 2 to 0.7 nm between each ceramide headgroup and the side chain of E73^−^, thereby effectively preventing ceramide from directly contacting this residue (**Fig. 2a, b**). When applying a restriction radius of 2 nm, HKI-N binding to VDAC1 was largely unaffected by C_16:0_ ceramide. However, when narrowing the restriction radius to 1.5 or below, complex assembly was strongly impaired (**Fig. 2c, d**), even though under these conditions, C_16:0_ ceramide had only access to the area immediately adjacent to the bilayer-facing E73^−^ but not to the residue itself (**Fig. 2b**). This suggests that C_16:0_ ceramide disrupts HKI binding to VDAC1 by altering the properties of the membrane region vicinal to the bilayer-facing Glu, rather than through direct competition for binding to it. This indirect mechanism is consistent with the aforementioned interference of ceramide on the formation of VDAC–HKI complexes, whereby preformed complexes are left unaffected.

### Ceramide reduces cytosolic leaflet thinning near the bilayer-facing Glu

VDACs cause a global membrane thinning around the entirety of their perimeter due to a mismatch of their hydrophobic thickness (∼2.4 nm) with that of the surrounding membrane bilayer (∼4 nm)^25^. Superimposed on this overall mismatch, VDAC1 and VDAC2 harbor a uniquely positioned set of polar residues that impose a pronounced thinning of the cytosolic leaflet near the charged Glu. We previously demonstrated that substitution of Leu for the key polar residues Ser101 and Thr77 in VDAC1 neutralized cytosolic leaflet thinning and impaired HKI binding^22^. Consequently, we considered that C_16:0_ ceramide may perturb VDAC-HKI complex assembly by modulating the thickness and compressibility of the cytosolic leaflet near the charged Glu. To test this, we simulated VDAC1 and VDAC2 with charged Glu residues in a POPC:SAPC (90:10 mol%) bilayer to which we added different concentrations of C_16:0_ ceramide at the expense of POPC. A homogeneous PC lipid background was used to facilitate bilayer thickness measurements via lipid phosphate groups because such measurements are less reliable when using the heterogeneous OMM-mimicking membrane model. Polyunsaturated SAPC was included to keep the POPC membrane in a fluid state, since in its absence, C_16:0_ ceramide causes a pure POPC bilayer to transition to the gel phase (**Suppl. Movie 1**), in line with known behavior^27^. When added to VDAC1 or VDAC2 simulated in a POPC:SAPC bilayer, C_16:0_ ceramide in each caused a gradual, concentration-dependent neutralization of cytosolic leaflet thinning near the charged Glu (**Fig. 3a-c**). C_16:0_ ceramide also increased the global thickness of the lipid bilayer both immediately around the channel perimeter and further away from the channel. However, C_16:0_ ceramide’s thickening properties were most pronounced in the cytosolic leaflet in the region immediately adjacent to the charged Glu (**Fig. 3c**). Importantly and conversely, no changes in bilayer thickness were observed when simulations were carried out in the presence of C_16:0_ sphingomyelin. Collectively, these results indicate that the thickness of the cytosolic leaflet near the charged Glu plays a decisive role in HKI-VDAC binding and is particularly susceptible to the membrane rigidifying properties of ceramides. Consequently, we envision that C_16:0_ ceramide perturbs HKI binding to VDACs by causing a thickening of the cytosolic leaflet in this area, thereby congesting the polar pathway used by HKI to gain access to the critical, membrane-buried Glu.

**Figure 3.**
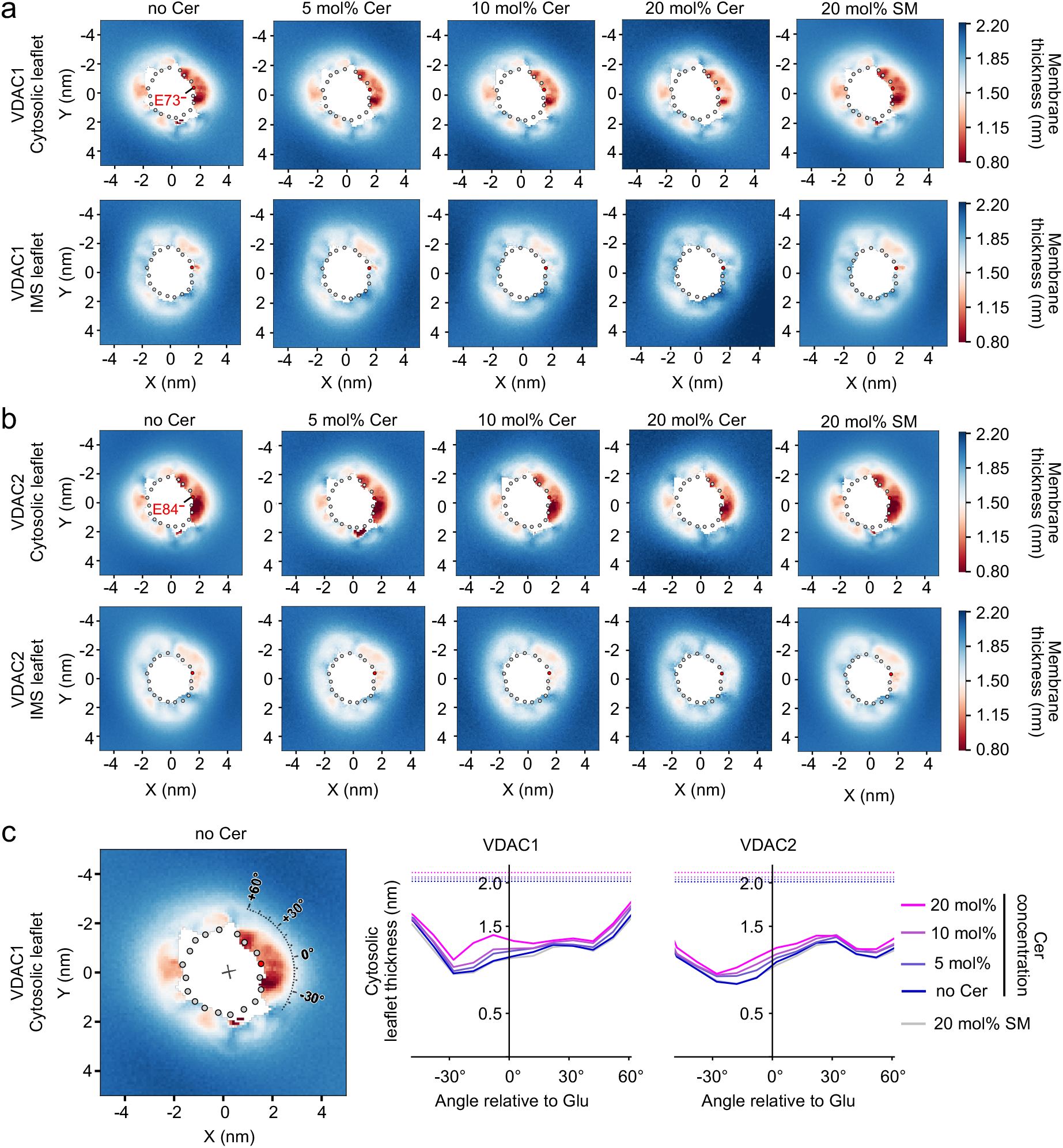
Ceramides reduce cytosolic leaflet thinning near the bilayer-facing Glu. **(a)** Leaflet-specific membrane thinning heatmaps of VDAC1 with a charged E73 simulated in a bilayer comprising POPC:SAPC supplemented with the indicated amount of C_16:0_ ceramide or C_16:0_ sphingomyelin. The amount of SAPC was kept constant at 10 mol% to keep the membrane bilayer fluid. The position of E73 is marked by a *red* dot. **(b)** Leaflet-specific membrane thinning heatmaps of VDAC2 with a charged E84 in simulations as in (a). The position of E84 is marked by a *red* dot. **(c)** Average cytosolic leaflet thickness measured along an arc with a radius of 2.5 nm from the barrel center of VDAC1 or VDAC2 as a function of the angle relative to the bilayer-facing Glu (depicted as an overlay in the leftmost plot). Measurements were from simulations of VDAC1 as in (a) or VDAC2 as in (b). Dotted lines mark the average cytosolic leaflet thickness outside of a 2.5 nm radius away from the barrel center.

### VDAC1 and VDAC2 are inherently tilted in the membrane bilayer

A previously overlooked feature of VDAC channels is their persistent tilt in the bilayer. This feature became apparent after we constructed a vector that was aligned with the β-barrel direction to assess the native tilt of both VDAC1 and VDAC2 in an OMM-mimicking bilayer, measured as the rotation away from verticality along the center→E73 axis or perpendicularly to it (**Fig. 4a**). Using this approach, we found that both channels are tilted in the opposite direction of the membrane-buried Glu, with VDAC2 displaying a steeper tilt, with an average deviation of 7.84° from the vertical, compared to 4.76° for VDAC1 (**Fig. 4b, c**). To explore the mechanism that drives VDAC tilting, we simulated VDAC1 and VDAC2 in a 100 mol% POPC bilayer under conditions whereby the inherent channel tilt was either left unaffected or restrained to keep it around 0°. Restraining the channel tilt caused a sharp increase in membrane defects in the cytosolic leaflet around the bilayer-facing Glu, a phenomenon that was most pronounced in simulations of the more tilted VDAC2 (**Fig. 4d**). Neutralizing channel tilt in this way affected the membrane organization on all sides of the protein, with new membrane defects emerging on the diagonally opposite side of the region where the cytosolic leaflet became thinner (**Fig. 4d, e**). These results reinforced the notion that tilting is an inherent feature of VDACs. As VDAC tilting effectively results in an enhanced cytosolic exposure of the membrane-buried Glu (**Fig. 4b**), we wondered whether constraining channel tilt toward 0° would affect HKI binding. However, neutralizing tilt had no (in case of VDAC1) or only a minor impact on HKI binding (in case of VDAC2; **Suppl. Fig. 3a-c**). This indicated that tilt is not a critical factor in HKI-VDAC complex formation, presumably because the deepening of membrane defects induced by restricting tilt keeps the critical membrane-buried Glu accessible for HKI binding.

**Figure 4.**
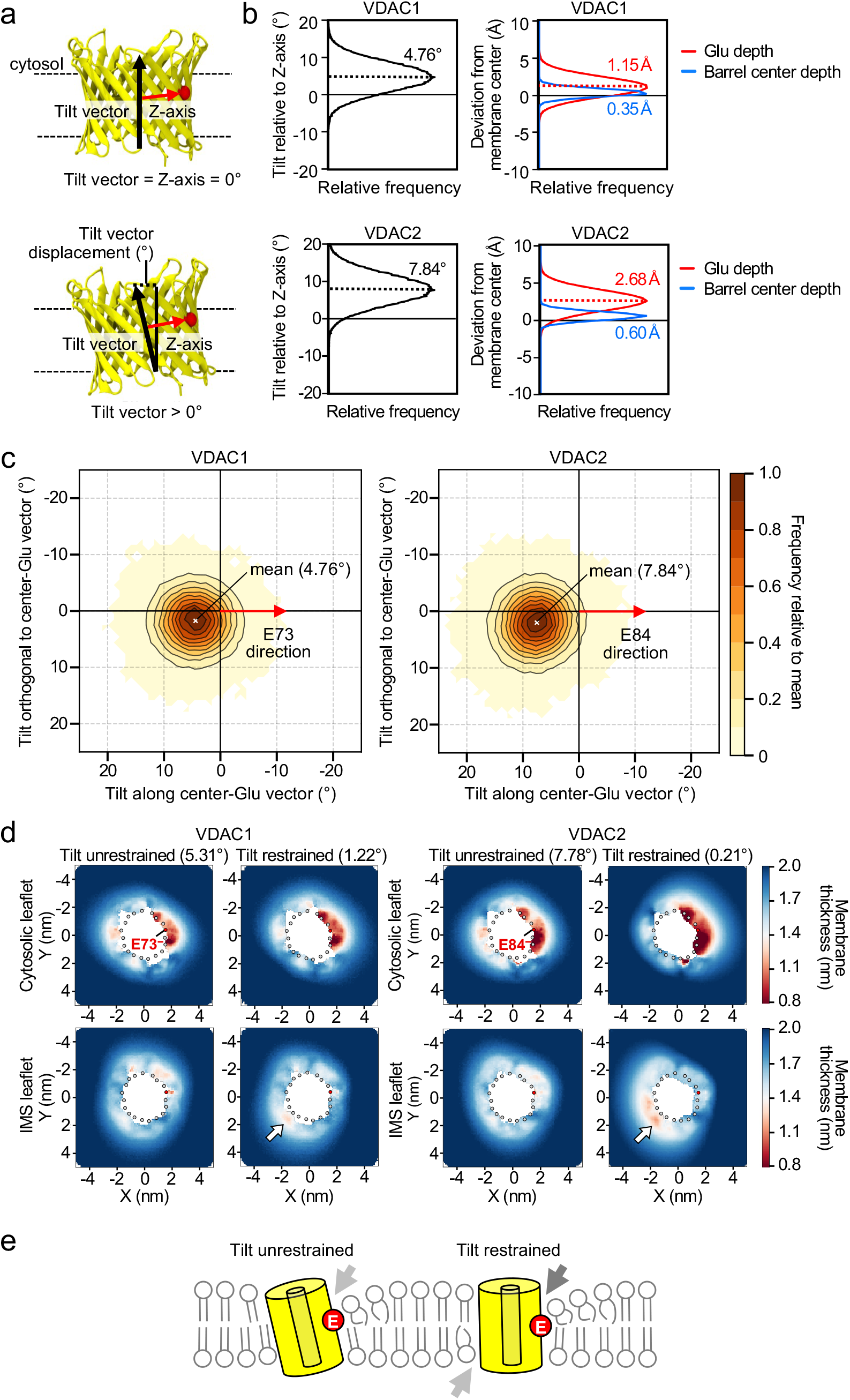
VDAC1 and VDAC2 are inherently tilted in the membrane bilayer. **(a)** Schematic of VDAC tilt measurements. A tilt vector was defined to correspond to the cylindrical axis of the VDAC barrel (see the Methods section for details on tilt vector construction). The rotation of this tilt vector was measured either as the angle to the z-axis (overall tilt) or discriminated around reference axes in the xy-plane (center→Glu axis or perpendicularly to it). The position of the charged, bilayer-facing Glu residue is marked by a *red* sphere. **(b)** Distribution of VDAC overall tilt (*left*) and membrane depth, including of the bilayer-facing Glu (*right*). VDAC channels were simulated in an OMM-mimicking bilayer with a deprotonated Glu. The depths, in z, of the barrel center and bilayer-facing Glu were measured relative to the geometric center of the membrane bilayer. Positive values indicate a displacement towards the cytosolic leaflet. All distributions are plotted normalized to their maximum value. **(c)** Tilt plots of VDAC1 and VDAC2 simulated as in (b). Distributions are plotted normalized to their maximum. The graph is plotted such that the x-axis corresponds to the vector between the barrel center and bilayer-facing Glu (the direction towards the Glu is indicated by an arrow). Positive values in x indicate tilting in the direction opposite of the bilayer-facing Glu. Note that, to conform to the convention where positive angles represent anti-clockwise rotations, values in x increase from right to left. **(d)** Leaflet-specific membrane thinning graphs of VDAC1 and VDAC2 channels simulated in a POPC bilayer with or without tilt restriction. Indicated are the mean tilt values of each channel relative to the z-axis over the course of the simulation. White arrows indicate areas of leaflet-specific membrane thinning emerging upon tilt restriction. **(e)** Schematic of how VDAC channels influence the membrane bilayer organization around the channel wall with or without tilt restriction.

### Polar residues adjacent to the membrane-buried Glu are key determinants of VDAC tilting

As VDAC1 and VDAC2 are tilted in the opposite direction of the membrane-buried Glu where an adjacent set of polar residues cause a pronounced thinning of the cytosolic leaflet, we considered that tilt might serve to minimize the energetically unfavorable hydrophobic mismatch that VDACs face in this region. Consistent with this notion, we observed that restraining channel tilt significantly amplified the membrane defects in the Glu-surrounding region (**Fig. 4d, e**). To better characterize the mechanism responsible for VDAC tilting, we analyzed the influence of the membrane-buried Glu and adjacent polar channel residues on the mean tilt of VDACs simulated in an OMM-mimicking bilayer. Protonation of the membrane-buried Glu or its replacement with Gln had only a modest neutralizing effect on channel tilt. Substitution of Leu for the key polar residues Ser101 and Thr77 in VDAC1, on the other hand, caused a more pronounced decrease in channel tilt, namely from ∼4.76° to ∼3.09° (**Fig. 5a**). Likewise, substitution of Leu for the corresponding polar residues Thr112 and Thr88 in VDAC2 resulted in a comparable decrease in channel tilt, from ∼7.84° to ∼5.09°. °. In all cases, decreases in tilt led to a concomitant deepening of the Glu in the membrane (**Suppl Fig. 4**). These results suggested that the hydrophobic mismatch imposed by polar residues on the cytosolic leaflet adjacent to the bilayer-facing Glu serves as a major driver of VDAC tilting, with the protonation state of the Glu residue playing a relatively minor role. To further validate this model, we next simulated wildtype and polar residue mutant variants of VDAC1 and VDAC2 under tilt-restrained and unrestrained conditions in a 100 mol% POPC bilayer. Under unrestrained conditions, the membrane defects in the Glu-adjacent region of VDAC1 and VDAC2 were either abolished or strongly reduced in the polar residue mutants VDAC1^S101L/T77L^ and VDAC2^T112L/T88L^, respectively (**Fig. 5b, c**). This was accompanied by a reduction in channel tilt from ∼5.31° to ∼3.39° for VDAC1 and from ∼7.78° to ∼7.08° for VDAC2. Strikingly, restraining the tilt of the polar residue mutant channels no longer led to a sharp increase in membrane defects in the cytosolic leaflet around the bilayer-facing Glu (**Fig. 5b, c**), as had previously been observed for the wild type channels (**Fig. 4d**). Collectively, these data indicate that the polar residues adjacent to the bilayer-facing Glu are key determinants of VDAC tilting.

**Figure 5.**
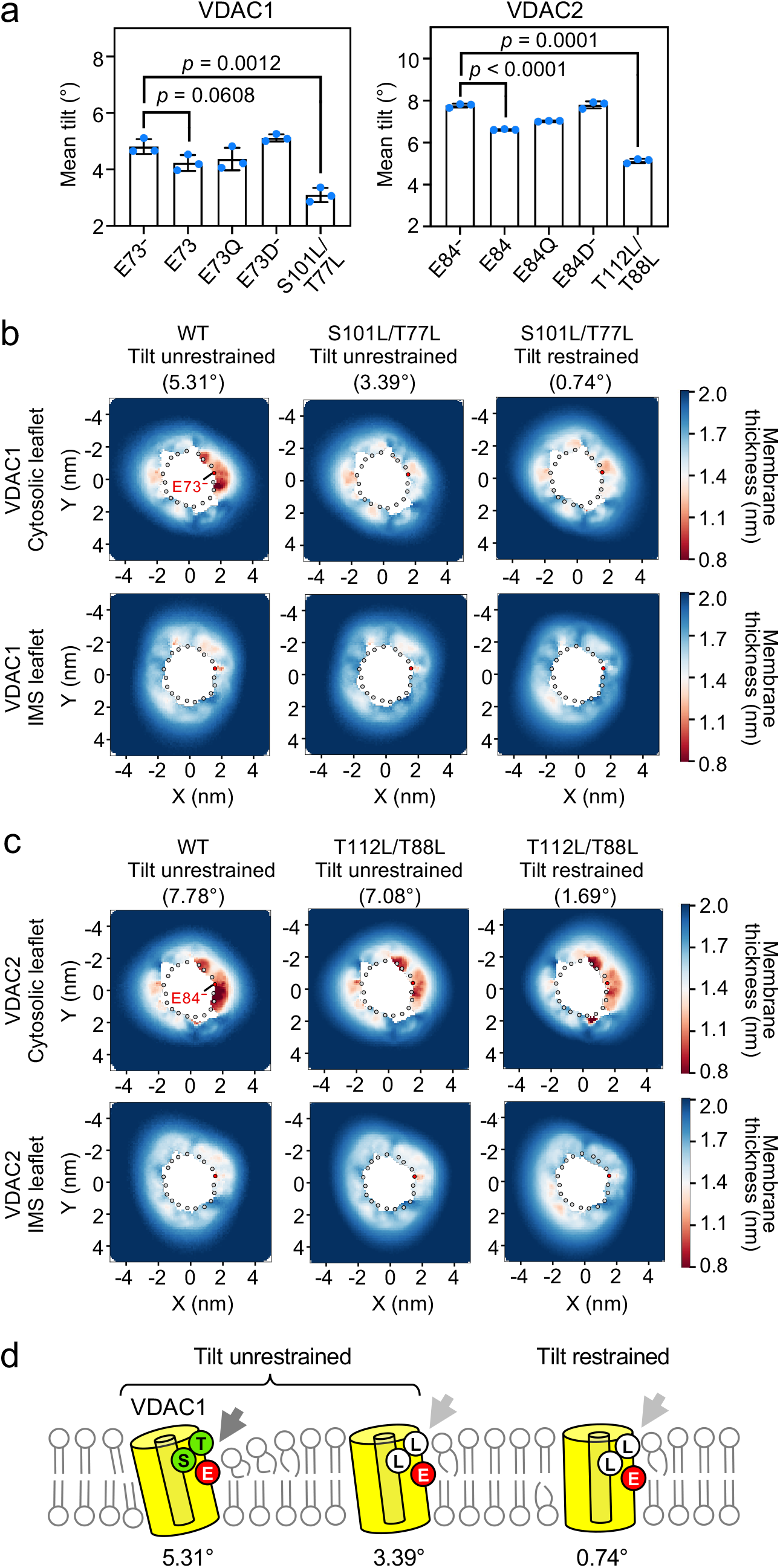
Polar residues adjacent to the bilayer-facing Glu contribute to VDAC tilting and cytosolic leaflet thinning. **(a)** Mean tilt values of wildtype and mutant VDAC channels simulated in an OMM-mimicking bilayer. VDAC1^S101L/T77L^ and VDAC2^T112L/T88L^ mutants were simulated with a deprotonated bilayer-facing Glu. Data are means ± SD, n = 3 per condition. *p* values were calculated by unpaired two-tailed *t* test. **(b)** Leaflet thinning graphs of VDAC1 and VDAC1^S101L/T77L^ simulated in a POPC bilayer with or without tilt restriction. Indicated are the mean tilt values of VDAC1 relative to the z-axis over the course of the simulation. **(c)** Leaflet thinning graphs of VDAC2 and VDAC2^T112L/T88L^ simulated in a POPC bilayer as in (b). Indicated are the mean tilt values of VDAC2 relative to the z-axis over the course of the simulation. **(d)** Schematic of how polar residues S101 and T77 in VDAC1 affect the membrane bilayer around the channel wall.

### HKI and ceramide are modulators of VDAC tilting with antagonistic effects

As C_16:0_ ceramide partially neutralized the cytosolic leaflet thinning mediated by polar residues near the bilayer-facing Glu of VDACs (**Fig. 3a-c**), we next investigated its potential impact on channel tilt. To this end, we simulated VDAC1 and VDAC2 in an OMM-mimicking bilayer containing different concentrations of C_16:0_ ceramide. This revealed that C_16:0_ ceramide caused a concentration-dependent decrease in the tilt of both VDAC1 and VDAC2 (**Fig. 6a**). In contrast, C_16:0_ sphingomyelin had no effect on channel tilt. To investigate whether HKI binding influenced channel tilt, we next simulated HKI-N with VDAC1 or VDAC2 in an OMM-mimicking bilayer and measured channel tilt in all frames in which HKI-N Met1 was in direct contact with the bilayer-facing Glu. This revealed that HKI binding to VDAC1 caused a minor but significant increase in channel tilt while HKI binding to VDAC2 had no impact (**Fig. 6a**). These results mirror the impact of C_16:0_ ceramide and HKI binding on the cytosolic exposure of the membrane-buried Glu (**Suppl. Fig. 5**). From this we conclude that C_16:0_ ceramide and HKI act as modulators of VDAC tilting, with ceramides neutralizing channel tilt and HKI binding having the opposite effect.

**Figure 6.**
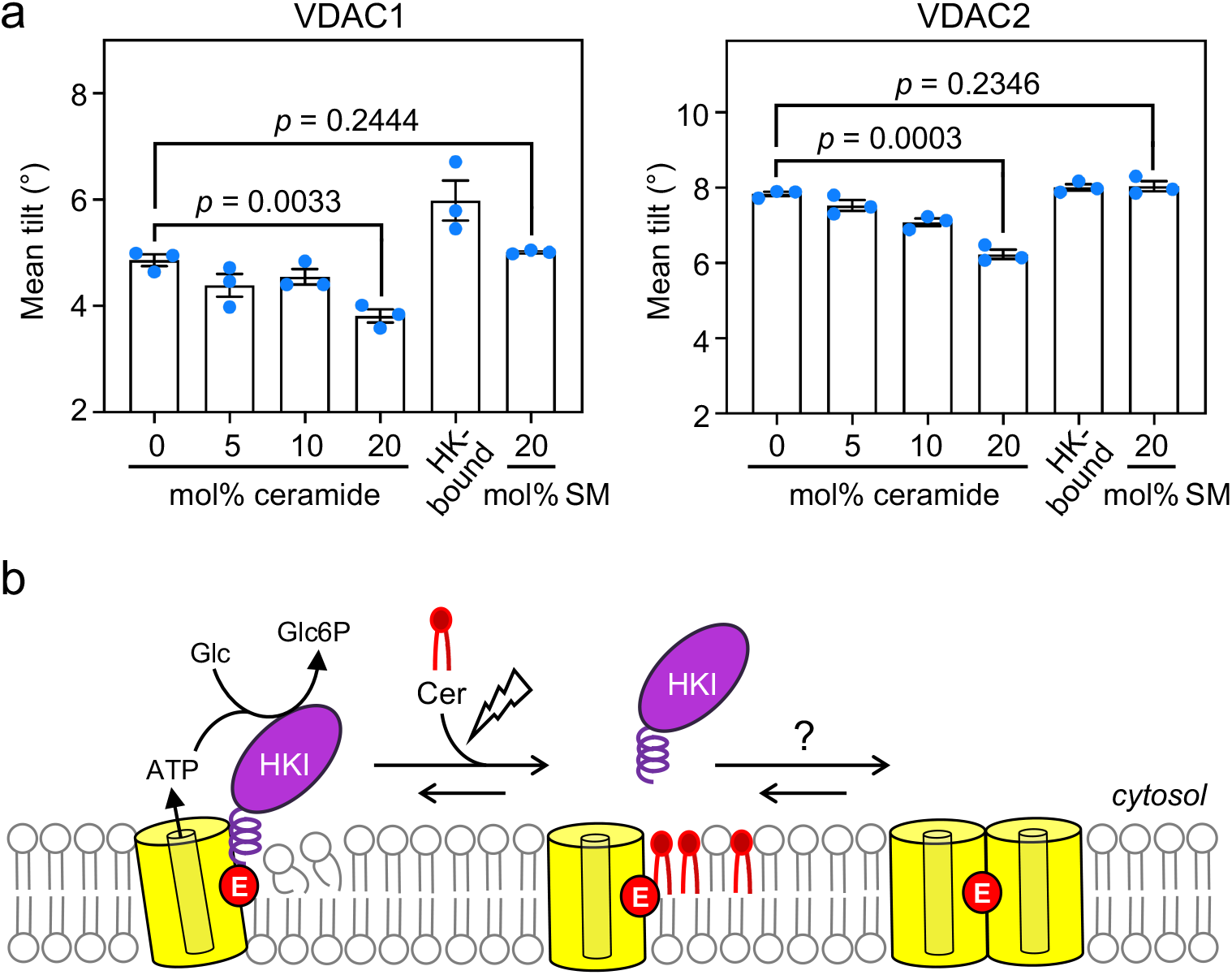
HKI and ceramides are antagonistic modulators of VDAC tilting. **(a)** Mean tilt values of VDAC1 and VDAC2 with a charged Glu simulated in an OMM-mimicking bilayer containing the indicated amount of C_16:0_ ceramide or C_16:0_ sphingomyelin. Mean tilt values of VDAC1 and VDAC2 channels with bound HKI-N were calculated from the simulations that yielded Fig. 1c and 1e. Data are means ± SD, n = 3 per condition. *p* values were calculated by unpaired two-tailed *t* test. **(b)** Cartoon illustrating how ceramides disrupt HKI-VDAC complex formation and modulate VDAC channel tilting. The lightening symbol marks an apoptotic stimulus that causes a rise in mitochondrial ceramide levels.

## DISCUSSION

Ceramide binding to VDACs promotes mitochondrial apoptosis, but the underlying mechanism is not clear. We previously showed that ceramide binding relies on a charged, bilayer-facing Glu that also controls the assembly of pro-survival complexes between VDACs and hexokinase HKI ^14,22^. Consequently, we postulated that ceramides may initiate mitochondrial apoptosis by displacing HKI from VDACs. Using CG-MD simulations, we here show that ceramides perturb HKI-VDAC complex assembly in a concentration dependent manner. However, this does not occur via direct competition between ceramides and HKI for binding to the crucial Glu. Rather, we find that ceramides selectively congest a polar pathway used by HKI to gain access to the membrane-buried Glu, thereby interfering with efficient VDAC-HKI complex assembly. Additionally, we observed that VDACs are inherently tilted in the membrane bilayer and identified polar channel residues adjacent to the membrane-buried Glu as key determinants of channel tilting. Our finding that ceramides and HKI antagonistically influence VDAC tilting provides new mechanistic insight into how VDAC channels participate in ceramide-mediated apoptosis.

HKI contains an 18-residue long *N*-terminal amphipathic α-helix, HKI-N, which is both essential and sufficient to mediate binding of the enzyme to VDAC1 and VDAC2^28,29^. Complex assembly relies on intimate contacts between HKI-N and the charged Glu on the outer wall of both channel isomers. Protonation of this residue blocks complex formation in simulations while transient acidification of the cytosol causes a reversable release of HKI-N from mitochondria^22^. Membrane insertion of HKI-N occurs adjacent of the charged Glu where a pair of polar channel residues causes a marked thinning of the cytosolic leaflet, creating a funnel that serves as low-energy passageway for HKI to reach the bilayer-facing Glu and form a stable complex. Consistent with this idea, we found that disrupting the membrane-thinning capacity of VDAC1 by mutation of this pair of polar residues significantly impaired VDAC1-HKI complex assembly both in simulations and in cells^22^. Here we show that when ceramides are restricted in such a way that they cannot interact with the bilayer-facing Glu but still have access to the funnel, the assembly of the HKI-VDAC complex was impeded to the same extent as under unrestricted conditions. This strongly suggests that disruption of HKI-VDAC complex assembly by ceramides is not due to their direct competition with HKI-N for binding to the bilayer-facing Glu. Instead, we found that ceramides that populate the funnel adjacent to the bilayer-facing Glu caused a marked thickening of the cytosolic leaflet. Unlike ceramides, sphingomyelin had no influence on leaflet thickness and did not impair HKI-VDAC binding. Ceramides exhibit a number of properties not shared by almost any other membrane lipid, including an extremely hydrophobic saturated backbone and a small headgroup that is only moderately polar^26^. These features explain a tendency of ceramides to intercalate between phospholipid acyl chains, which drives lipid packing and increased bilayer thickness and stiffness^30^. Our MD simulations indicate that the membrane area proximal to the membrane-buried Glu and polar patch on the outer channel wall of VDACs is particularly susceptible to changes in the lipid environment. We propose that the ceramide-induced thickening and rigidification of the cytosolic leaflet in this area congests the gateway for HKI to bind VDACs, blocking off access of the enzyme to the critical membrane-buried Glu.

The impact of ceramides on leaflet thickness and HKI-VDAC binding in simulations became apparent at a relative high ceramide concentration (10 mol%). While ceramides are usually found in smaller amounts (<1 mol%) in cellular membranes, their local concentration may increase by tenfold and even more under stress or apoptosis conditions^6,31,32^. Previous work revealed that mitochondrial ceramide generation is obligate for radiation-induced apoptosis, involving activation of ceramide synthases in mitochondria-associated membranes (MAM) and an apparent transfer of newly synthesized ceramide to the OMM^5,13,33^. This resulted in the formation of mitochondrial ceramide-rich macrodomains in which VDACs, Bax and Bak were found concentrated^13^. Under these conditions, the concentration of ceramides in the OMM may locally reach levels close to those used in this work.

A remarkable finding of our study was that VDAC1, and notably VDAC2, are inherently tilted in the membrane. Both channels are tilted in the opposite direction of the bilayer-facing Glu and surrounding polar region, suggesting that VDAC tilting is caused by the pronounced hydrophobic mismatch that these channels experience in this area. This notion is supported by the following observations. First, we found that VDAC2, which exerts a higher degree of leaflet thinning in the polar region than VDAC1, is also nearly twice as tilted. Second, substitution of Leu for the polar residues responsible for leaflet thinning in the Glu-adjacent region reduced the overall tilt of each channel isomer by almost half. Third, ceramides caused a concentration-dependent neutralization of channel tilting that correlated well with the extent to which these lipids imposed a thickening of the cytosolic leaflet near the bilayer-facing Glu. In contrast, sphingomyelin did not affect leaflet thickness in this area and also did not influence channel tilting. Collectively, these results indicate that the hydrophobic mismatch between VDACs and the lipid bilayer in the polar region surrounding the membrane-buried Glu drives channel tilting, presumably to minimize the unfavorable exposure of this region to the hydrophobic membrane core.

A persistent tilt in the membrane bilayer is a previously overlooked feature of VDAC channels. This raises the question how VDAC tilting may impact channel function. Although VDAC monomers can govern ion and metabolite fluxes across the OMM, VDAC oligomerization has been implicated in a variety of physiological processes, including apoptosis and mitochondrial DNA release^34,35^. However, the molecular drivers of VDAC oligomerization remain unclear. There is compelling evidence that VDAC1 oligomerization is influenced by the membrane lipid composition^25^. Moreover, previous work revealed that protonation of the membrane-buried Glu in VDAC1 promotes the formation of channel dimers, with the Glu positioned at the dimer interface^36^. In this context, it is important to note that protonation of this glutamate residue reduced channel tilting to a similar extent as a high ceramide concentration, especially in the case of VDAC2 (**Fig. 5a, Fig. 6a**). It is plausible that VDAC tilting impedes channel dimerization. This would keep the bilayer-facing Glu accessible to HKI to form a heterodimeric complex that shields a VDAC interface required for channel oligomerization as part of an anti-apoptotic response. Our present findings indicate that ceramides may reverse these events and facilitate the formation of pro-apoptotic VDAC oligomers by neutralizing channel tilt and preventing HKI-VDAC complex assembly (**Fig. 6b**). This model provides a useful guide for futures studies aimed at unraveling the mechanism by which ceramides exert their widely acclaimed anti-neoplastic activities.

## METHODS

### Computational models and structures

Computational models and structures were used according to Bieker et al.^22^ with some modifications. Briefly, for all CG-MD simulations, an NMR structure of human VDAC1 (PDB:6TIQ) and the first 18 residues of the *N*-terminal helix of rat HKI-structure (PDB:1BG3), which is identical in sequence to the non-resolved structure of the human HKI *N*-terminus, were used. For VDAC2, we truncated the first eleven residues to achieve the same sequence length as VDAC1 and then mutated the structure of human VDAC1 onto the human VDAC2 sequence. An RMSD of 2.03 Å between the used human VDAC1 β-barrel backbone and an available zebrafish VDAC2 structure (PDB:4BUM) strongly supports our assumption of identical secondary structure between the two proteins. The martinize2 script^37^ was used to coarse-grain all proteins. We protonated two protein sites to more closely mimic physiological conditions: Firstly, we protonated the HKI-N *C*-Terminus to mimic a continuation of the protein. Secondly, we protonated a barrel-lumen facing aspartate (D100 in VDAC1 and D111 in VDAC2), as it was falling just inside the electrostatic distance cutoff range of HKI-N, when it was in contact with Glu73/84, while two adjacent lysines fell just outside the electrostatic distance cutoff, possibly overrepresenting the aspartate’s negative charge.

VDAC1 was embedded into an OMM-mimicking bilayer using the insane-script^38^. The membrane composition was based on Horvath and Daum^39^, with the cytosolic leaflet consisting of POPC/POPE/SAPI/cholesterol (45/33.5/5/16.5, mol%) and the IMS-leaflet consisting of POPC/POPE/SAPI/cholesterol (52.5/14/19/14.5, mol%). When C16:0 ceramide or C16:0 sphingomyelin was added to these membrane systems, all other lipids were reduced accordingly to maintain their initial relative composition ratios. For membrane thickness and defects measurements, we used the insane script to either embed VDAC1 in a 100 mol % POPC membrane to ensure leaflet symmetry or a symmetric POPC/SAPC (90/10, mol%) lipid mixture to additionally keep the membrane in a fluid state at high ceramide concentrations. When C16:0 ceramide or C16:0 sphingomyelin was added to these membranes, we reduced the concentration of POPC accordingly and kept SAPC at a stable concentration.

In case of all VDAC2 simulations, the VDAC2 backbone particles were aligned onto the membrane-embedded VDAC1 backbone particles using the MDAnalysis^40^ and MDreader (https://github.com/mnmelo/MDreader) Python packages. This ensured a similar tilt and starting position between the two proteins. Approximately 150 mM NaCl was added to all systems, with an excess of Na+ to reach charge neutrality.

### MD simulation settings

All unrestrained simulation systems were set up and simulated as in Bieker et al.^22^, with modifications for restrained systems. Briefly, we simulated all systems using the CG Martini 3 force field^41^ with updated lipid models^42^ and restricted the secondary structure of all proteins using an elastic network approach^43^. All simulations were carried out using GROMACS versions 2021^44^. For all conditions, we first employed a steepest-descent energy minimization then, in triplicate, equilibrated temperature and pressure and simulated production runs.

For equilibrations with HKI-N, the peptide was subjected to a flat-bottom harmonic restraint in z and a harmonic restraint in x and y to both ensure close proximity to the bilayer and to prevent any movement away from its initial placement at the box edge. These restraints were lifted after a membrane-adsorbed state of the HKI-N peptide was confirmed after equilibration. During the production run, a repulsive harmonic potential acting only on the peptide was put in place with onset 1.5 nm away from the z=0 position. This had no effect on the peptide when membrane-adsorbed, but prevented it from crossing the system’s periodicity in z and contacting the opposite leaflet if it left the membrane.

For simulations where intermolecular distances or tilts were restrained, we employed the PLUMED plugin for molecular dynamics simulations^45^. For distance restraints between the ceramide headgroup beads and the membrane-buried Glu73 sidechain bead, we employed a constant harmonic distance restraint in a sphere around Glu73 with a force constant of 700 kJ/mol/nm^2^ at indicated restraint distances. For VDAC tilt measurement and restraining the follow procedure was employed: we defined two rings of backbone beads from the β-sheet portion of the VDAC barrel: one closest to the IMS leaflet and the other one closest to the cytosolic leaflet. Each ring consisted of 19 backbone beads, one per each β-sheet of the barrel. We then defined points *c*_1_ and *c*_2_ at the geometric center of the IMS-leaflet or cytosolic leaflet rings, respectively. The VDAC barrel center, *c*_VDAC_, was defined as the midpoint between *c*_1_ and *c*_2_. Another reference point was the backbone bead of the membrane-buried Glu backbone bead (*c*_Glu_). With these points we then defined the barrel vector as 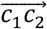, the first reference vector as 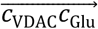, and a second reference vector, 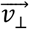, perpendicular to both 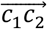, and 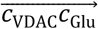, (computed as the cross product 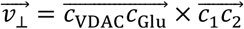. Tilts were measured as the torsion of the barrel vector away from the z-axis either around the 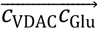 or the 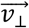, vectors (always assuming *c*_VDAC_ as the pivot point). To restrain VDAC tilt closer to 0°, we employed constant forces on the barrel vector rotation around both 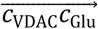 and 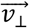, with values dependent on the VDAC variant or mutant. To adjust VDAC1 tilt so it approached VDAC2 tilt, we employed a constant force 25 of kJ/mol/rad for rotation around the 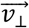 vector. An overall tilt measurement was also defined over a single rotation axis 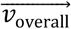 for each system, the time average of the barrel vectors 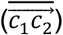 was computed, and 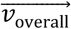 defined as the cross product between the z axis and it 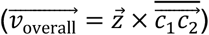. This definition allowed computing tilt rotations in the [−*π, π*[ domain, which is a relevant distinction to simply measuring the angle between vectors, particularly when approaching collinearity. The Data Availability section links to a repository of all used plumed files, starting configurations and indices for the respective restrained conditions.

### Simulation analysis

Simulation analyses of HKI-N–VDAC contacts, ceramide headgroup occupancy and membrane defects were carried out according to Bieker et al.^22^. Briefly, for HKI-N binding to VDACs, we analyzed contacts between the proteins’ particles within a 0.6 nm cutoff, grouped into contacts per residue (in which residues were considered in contact if any of their beads were in contact). These were plotted as either HKI-N Met1 contacts to any VDAC residue or as any HKI-N residue to Glu73/84, respectively. Contact intensity is shown as the fraction of total simulation time for which the contact was established. To define contact events and measure their lifetimes, we set Met1-Glu distance cutoffs at 0.6 and 1.0 nm; a contact event was defined as a continuous range of frames in which the distance was always inside the 1.0 nm cutoff, as long as it reached below the 0.6 nm cutoff at some point in that interval. Contact durations were histogrammed into logarithmic bins, and plotted after weighing for the total contact duration per bin and normalizing for total simulated time. Average contact lifetimes were also calculated weighting for contact duration. For ceramide headgroup binding to VDACs, we considered a contact to be established, if any ceramide headgroup bead was in contact with any VDAC residue.

For ceramide headgroup and sphingomyelin phosphate occupancies, we used the VMD VolMap Tool^46^ to calculate the presence of a ceramide headgroup bead in a grid with a 0.2 nm spacing throughout the simulation box relative to the total simulation time. This leads to a possible occupancy range between 0% (never present) and 100% (always present). For membrane thickness heatmaps, we calculated the difference between the average z-position of all POPC backbone phosphates and the global membrane center of mass over an x/y-plane grid of 0.1 nm cell side. Values were plotted only if the phosphate count for the respective cell exceeded 30 counts over the entire trajectory.

To measure membrane thinning as a function of the position around VDACs, the average leaflet thickness values inside an arc with a radius of 2.5 nm from the barrel center and spanning approximately 110° in the Glu-adjacent area was determined and plotted. Barrel tilts were measured for each simulation frame. Angular distributions were plotted either as one-dimensional line distributions or as two-dimensional heatmaps normalized to their maximum.

For barrel and Glu depth distributions, the membrane center was defined as the midpoint between the centers of geometry of the phospholipid glycerols of each leaflet. The z position of the barrel center or the membrane-buried Glu were defined as in tilt restraint simulations, and measured relative to the membrane center. For tilt/depth measurements of the HKI bound state, the same methods were employed but analyzing only the simulation frames in which HKI-Met1 was in direct contact with the membrane-buried Glu, i.e., inside a 0.6 nm distance cutoff.

### Statistical analysis

The number of replicates as well as total simulation time of the combined replicates for each condition are given in the figure legends. Data are plotted either as cumulative data for each condition or as individual replicates with bar charts representing the mean and standard error of mean. *p* values were calculated by unpaired two-tailed t-test. For estimating the statistical error involved in the bound time fractions, simulations were divided in blocks of sufficient duration to be considered independent. This minimum block time was computed as the bound/unbound integrated auto-correlation time of each system.

## Acknowledgements

We gratefully acknowledge Ladislav Bartos and Robert Vácha (National Centre for Biomolecular Research, Masaryk University, Brno, Czech Republic) for providing the scripts for membrane thinning analysis. This work was supported by the Deutsche Forschungsgemeinschaft (HO3539/1-2 project no. 378148610 and HO 3539/2-1 project no. 448344643 to J. C. M. H.) and the FCT – Fundação para a Ciência e a Tecnologia I.P. (through MOSTMICRO-ITQB R&D Unit with projects UIDB/04612/2020 and LS4FUTURE Associated Laboratory with projects LA/P/0087/2020 and fellowship ITQBNOVA/FCT-Tenure/25/2025 to M. N. M.).

## Author Contributions

M. N. M. and J. C. M. H. designed the research with critical input from M. T.; M. T. performed all experiments with critical input from M. N. M.; M. N. M provided expertise for CG-MD simulations; M. N. M. and M. T. analyzed and interpreted the data, with critical input from J. C. M. H.; M. T. and J. C. M. H. wrote the manuscript with critical input from M. N. M.

## Data Availability

All data generated or analyzed in this study are included in the manuscript and supporting files. Source data with sample sizes, number of replicates, means, standard deviations, and calculated *p* values (where applicable) are provided in the Supplementary Data file.

## Code Availability

Models and code related to MD simulation preparation and analysis have been deposited in the Zenodo repository under doi: https://doi.org/10.5281/zenodo.19662948.

## REFERENCES

1. Chitkara, S. & Atilla-Gokcumen, G. E. Decoding ceramide function: how localization shapes cellular fate and how to study it. Trends in Biochemical Sciences vol. 50 Preprint at 10.1016/j.tibs.2025.01.007 (2025).

2. Patwardhan, G. A., Beverly, L. J. & Siskind, L. J. Sphingolipids and mitochondrial apoptosis. J. Bioenerg. Biomembr. 48, 153–168 (2016).

3. Bose, R. et al. Ceramide synthase mediates daunorubicin-induced apoptosis: An alternative mechanism for generating death signals. Cell https://doi.org/10.1016/0092-8674(95)90429-8 (1995) xdoi:10.1016/0092-8674(95)90429-8.

4. García-Ruiz, C. et al. Defective TNF-α-mediated hepatocellular apoptosis and liver damage in acidic sphingomyelinase knockout mice. Journal of Clinical Investigation 111, (2003).

5. Deng, X. et al. Ceramide Biogenesis Is Required for Radiation-Induced Apoptosis in the Germ Line of C. elegans. Science (1979). 322, 110–115 (2008).

6. Mesicek, J. et al. Ceramide synthases 2, 5, and 6 confer distinct roles in radiation-induced apoptosis in HeLa cells. Cell. Signal. 22, (2010).

7. Green, D. R. The Mitochondrial Pathway of Apoptosis Part I: MOMP and Beyond. Cold Spring Harb. Perspect. Biol. 14, (2022).

8. Birbes, H. et al. A mitochondrial pool of sphingomyelin is involved in TNFα-induced Bax translocation to mitochondria. Biochemical Journal 386, (2005).

9. Jain, A., Beutel, O., Ebell, K., Korneev, S. & Holthuis, J. C. M. Diverting CERT-mediated ceramide transport to mitochondria triggers Bax-dependent apoptosis. J. Cell Sci. 130, (2017).

10. Jain, A., Dadsena, S. & Holthuis, J. C. M. A switchable ceramide transfer protein for dissecting the mechanism of ceramide-induced mitochondrial apoptosis. FEBS Lett. https://doi.org/10.1002/1873-3468.13956 (2020) xdoi:10.1002/1873-3468.13956.

11. Schröer, C. et al. Mitochondria-specific photorelease of ceramide induces apoptosis. J. Lipid Res. 66, (2025).

12. Colombini, M. Ceramide channels and mitochondrial outer membrane permeability. Journal of Bioenergetics and Biomembranes Preprint at 10.1007/s10863-016-9646-z (2017).

13. Lee, H. et al. Mitochondrial ceramide-rich macrodomains functionalize bax upon irradiation. PLoS One https://doi.org/10.1371/journal.pone.0019783 (2011) doi:10.1371/journal.pone.0019783.

14. Dadsena, S. et al. Ceramides bind VDAC2 to trigger mitochondrial apoptosis. Nat. Commun. 10, (2019).

15. De Pinto, V. Renaissance of vdac: New insights on a protein family at the interface between mitochondria and cytosol. Biomolecules 11, (2021).

16. Jahn, H. et al. Phospholipids are imported into mitochondria by VDAC, a dimeric beta barrel scramblase. Nat. Commun. 14, (2023).

17. Chin, H. S. et al. VDAC2 enables BAX to mediate apoptosis and limit tumor development. Nat. Commun. 9, (2018).

18. Lauterwasser, J. et al. The porin VDAC2 is the mitochondrial platform for Bax retrotranslocation. Sci. Rep. 6, (2016).

19. Abu-Hamad, S., Zaid, H., Israelson, A., Nahon, E. & Shoshan-Barmatz, V. Hexokinase-I protection against apoptotic cell death is mediated via interaction with the voltage-dependent anion channel-1: Mapping the site of binding. Journal of Biological Chemistry 283, (2008).

20. Schindler, A. & Foley, E. Hexokinase 1 blocks apoptotic signals at the mitochondria. Cell. Signal. 25, (2013).

21. Lauterwasser, J. et al. Hexokinases inhibit death receptor-dependent apoptosis on the mitochondria. Proc. Natl. Acad. Sci. U. S. A. 118, (2021).

22. Bieker, S. et al. Hexokinase-I directly binds to a charged membrane-buried glutamate of mitochondrial VDAC1 and VDAC2. Commun. Biol. 8, 212 (2025).

23. Quach, C. H. T. et al. Mild alkalization acutely triggers the Warburg effect by enhancing hexokinase activity via voltage-dependent anion channel binding. PLoS One 11, (2016).

24. Shoshan-Barmatz, V., Ben-Hail, D., Admoni, L., Krelin, Y. & Tripathi, S. S. The mitochondrial voltage-dependent anion channel 1 in tumor cells. Biochimica et Biophysica Acta - Biomembranes vol. 1848 Preprint at 10.1016/j.bbamem.2014.10.040 (2015).

25. Lafargue, E. et al. Lipid composition of the membrane governs the oligomeric organization of VDAC1. Preprint at 10.1101/2024.06.26.597124 (2024).

26. Alonso, A. & Goñi, F. M. The Physical Properties of Ceramides in Membranes. Annu. Rev. Biophys. 47, 633–654 (2018).

27. Pinto, S. N., Silva, L. C., Futerman, A. H. & Prieto, M. Effect of ceramide structure on membrane biophysical properties: The role of acyl chain length and unsaturation. Biochim. Biophys. Acta Biomembr. 1808, (2011).

28. Gelb, B. D. et al. Targeting of hexokinase 1 to liver and hepatoma mitochondria. Proc. Natl. Acad. Sci. U. S. A. 89, (1992).

29. Ehsani-Zonouz, A., Golestani, A. & Nemat-Gorgani, M. Interaction of hexokinase with the outer mitochondrial membrane and a hydrophobic matrix. Mol. Cell. Biochem. 223, (2001).

30. García-Arribas, A. B., Busto, J. V., Alonso, A. & Goni, F. M. Ceramide and Cholesterol Effects on Phospholipid Bilayers under the AFM: Characterization of Complex Lipid Phases. Biophys. J. 108, (2015).

31. Wiesner, D. A. & Dawson, G. Staurosporine induces programmed cell death in embryonic neurons and activation of the ceramide pathway. J. Neurochem. 66, (1996).

32. García-Ruiz, C., Colell, A., Marí, M., Morales, A. & Fernández-Checa, J. C. Direct effect of ceramide on the mitochondrial electron transport chain leads to generation of reactive oxygen species: Role of mitochondrial glutathione. Journal of Biological Chemistry 272, (1997).

33. Stiban, J., Caputo, L. & Colombini, M. Ceramide synthesis in the endoplasmic reticulum can permeabilize mitochondria to proapoptotic proteins. J. Lipid Res. 49, (2008).

34. Keinan, N., Tyomkin, D. & Shoshan-Barmatz, V. Oligomerization of the Mitochondrial Protein Voltage-Dependent Anion Channel Is Coupled to the Induction of Apoptosis. Mol. Cell. Biol. 30, 5698–5709 (2010).

35. Kim, J. et al. VDAC oligomers form mitochondrial pores to release mtDNA fragments and promote lupus-like disease. Science (1979). 366, (2019).

36. Bergdoll, L. A. et al. Protonation state of glutamate 73 regulates the formation of a specific dimeric association of mVDAC1. Proc. Natl. Acad. Sci. U. S. A. 115, (2017).

37. Kroon, P. C. et al. Martinize2 and Vermouth provide a unified framework for molecular topology generation. Elife 12, (2025).

38. Wassenaar, T. A., Ingólfsson, H. I., Böckmann, R. A., Tieleman, D. P. & Marrink, S. J. Computational lipidomics with insane: A versatile tool for generating custom membranes for molecular simulations. J. Chem. Theory Comput. 11, (2015).

39. Horvath, S. E. & Daum, G. Lipids of mitochondria. Progress in Lipid Research vol. 52 Preprint at 10.1016/j.plipres.2013.07.002 (2013).

40. Gowers, R. et al. MDAnalysis: A Python Package for the Rapid Analysis of Molecular Dynamics Simulations. in Proceedings of the 15th Python in Science Conference (2016). doi:10.25080/majora-629e541a-00e.

41. Souza, P. C. T. et al. Martini 3: a general purpose force field for coarse-grained molecular dynamics. Nat. Methods 18, (2021).

42. Pedersen, K. B. et al. The Martini 3 Lipidome: Expanded and Refined Parameters Improve Lipid Phase Behavior. Preprint at 10.26434/chemrxiv-2024-8bjrr (2024).

43. Periole, X., Cavalli, M., Marrink, S. J. & Ceruso, M. A. Combining an elastic network with a coarse-grained molecular force field: Structure, dynamics, and intermolecular recognition. J. Chem. Theory Comput. 5, (2009).

44. Hess, B., Kutzner, C., Van Der Spoel, D. & Lindahl, E. GRGMACS 4: Algorithms for highly efficient, load-balanced, and scalable molecular simulation. J. Chem. Theory Comput. 4, (2008).

45. Bonomi, M. et al. PLUMED: A portable plugin for free-energy calculations with molecular dynamics. Comput. Phys. Commun. 180, (2009).

46. Humphrey, W., Dalke, A. & Schulten, K. VMD: Visual molecular dynamics. J. Mol. Graph. 14, (1996).

