## Supplementary Information for "Ceramides disrupt hexokinase HKI-VDAC complex assembly and modulate VDAC tilting"

**Affiliations:**

**This PDF file includes:**

Supplementary Figs. 1 to 5

Legend of Supplementary Movie 1

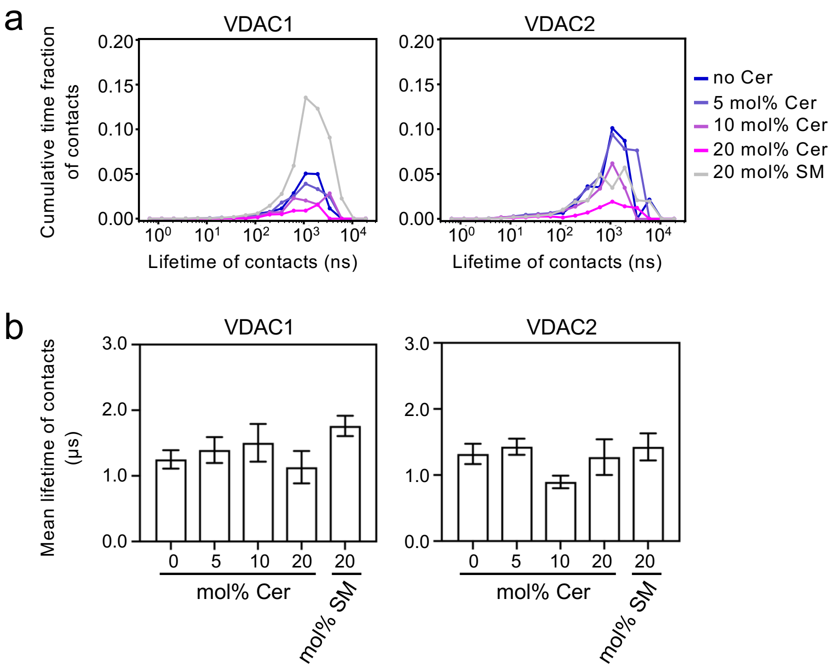


**Supplementary Figure 1. Distribution of contact lifetimes between HKI-Met1 and VDAC1-E73 or VDAC2-E84 in control or ceramide-containing bilayers.**

(**a**) Distribution of HKI-Met1–VDAC1-E73 or HKI-Met1–VDAC2-E84 binding events as a function of their lifetimes in control or ceramide-containing bilayers. Binding event lifetimes from all replicates were histogrammed over logarithmically-spaced bins, weighted by the summed lifetimes of the events in each bin and presented normalized by the total simulation time of each system's replicates.

(**b**) Mean binding lifetimes ± SEM of the same data as in (a). Supplementation of ceramide leads to a lower overall binding affinity of HKI for both VDAC1 and VDAC2, as evidenced by lower histogram values in (a), while similar average binding times are retained. This indicates that the reduced binding is a result of a lower on-rate, rather than shorter contact lifetimes — the converse would have entailed a distribution shift to lower lifetimes with increasing ceramide concentrations.

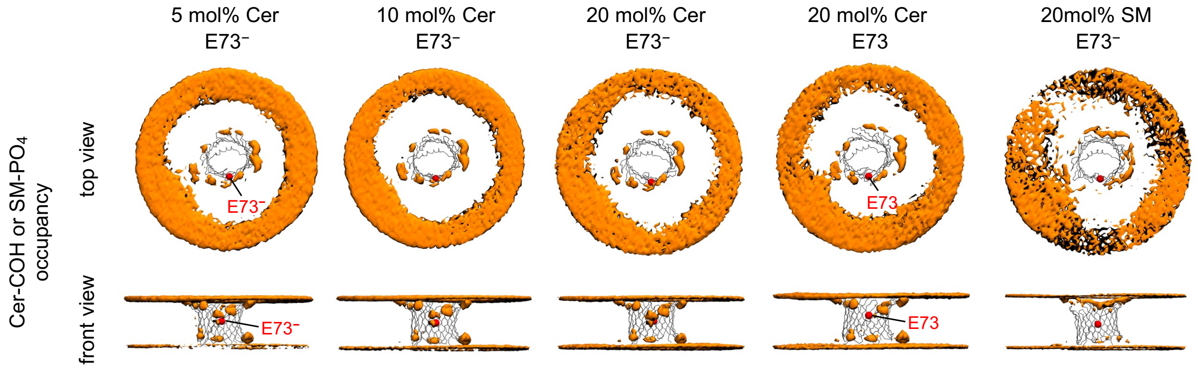


**Supplementary Figure 2. Ceramide distribution in the bilayer is consistent at different ceramide concentrations.**

Front and top view of calculated occupancies (orange) for the COH particle of $C_{16:0}$ ceramide (Cer) or the PO4 particle of $C_{16:0}$ sphingomyelin (SM). The position of the membrane buried Glu is indicated as a red sphere. Occupancy surfaces in the 5 mol% $C_{16:0}$ ceramide condition enclose volumes with an average occupancy of 0.1% or greater. The occupancy thresholds at higher $C_{16:0}$ ceramide or $C_{16:0}$ sphingomyelin concentrations were increased accordingly to account for higher density of sampled particles. This resulted in the following occupancy thresholds: 5 mol% Cer ≥ 0.1%; 10 mol% Cer ≥ 0.2%; 20 mol% Cer ≥ 0.4%; 20 mol% SM ≥ 0.4%. Overall, the same regions are visited, and in the same proportion, for the different ceramide concentrations simulated for VDAC with a deprotonated $E73^-$. Conversely and notably, when this residue is protonated the ceramide binding region directly adjacent to E73 is specifically abolished. Use of sphingomyelin instead of ceramide also yields no E73-adjacent occupied regions.

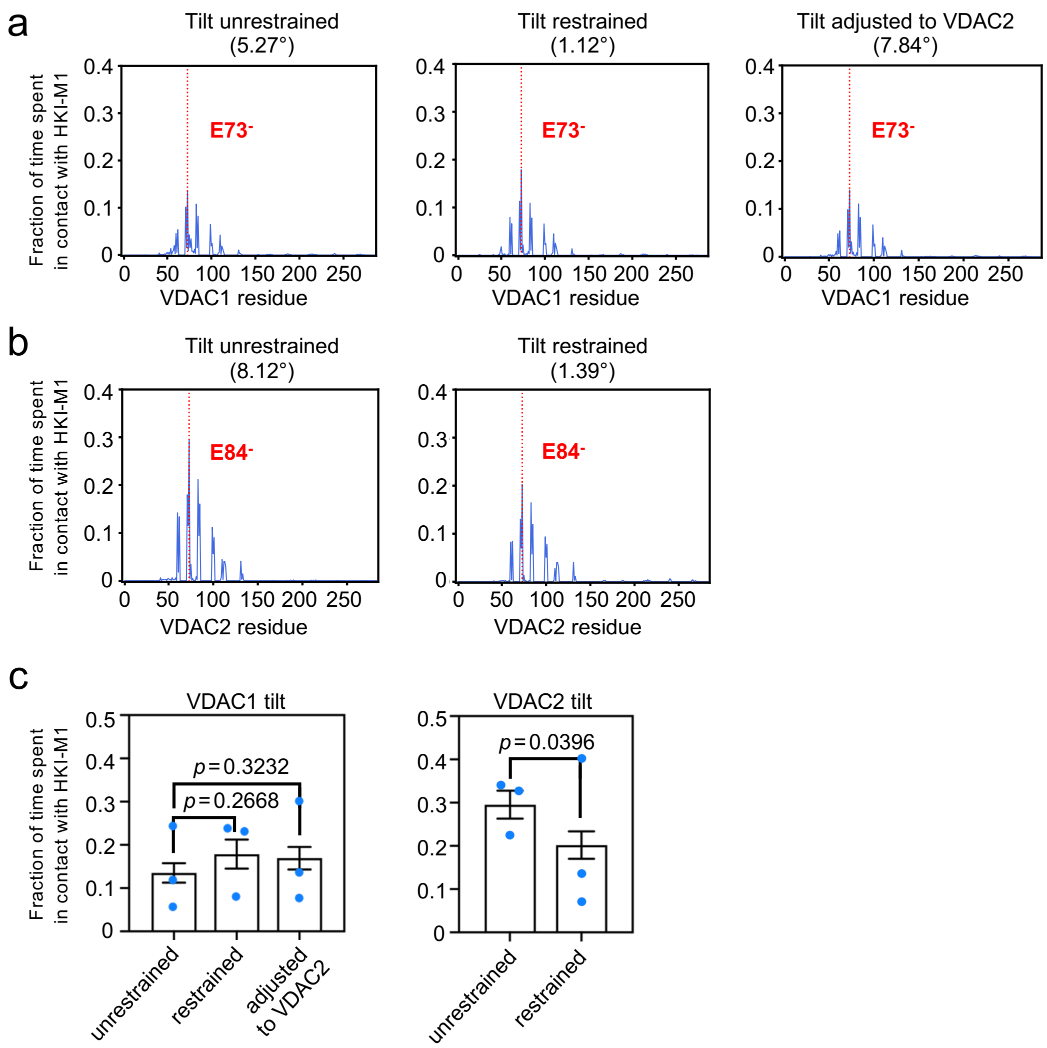


**Supplementary Figure 3. HKI binding to VDACs is not influenced by channel tilting.**

(**a**) Time fraction of contacts between HKI-Met1 and specific residues of VDAC1 with a deprotonated Glu73 under tilt unrestrained and tilt restrained conditions. Simulations were carried out in an OMM-mimicking bilayer. VDAC1 tilt was either neutralized or increased to mimic the inherent VDAC2 tilt. Shown are the combined data of three individual replicates per condition with a total simulation time between 215 μs and 233 μs per condition.

(**b**) Time fraction of contacts between HKI-Met1 and specific residues of VDAC2 with a deprotonated Glu84 under tilt unrestrained and tilt restrained conditions. Simulations were carried out in an OMM-mimicking bilayer. Shown are the combined data of three individual replicates per condition with a total simulation time between 225 μs and 230 μs per condition.

(**c**) Time fraction of contacts between HKI-Met1 and VDAC1 Glu73. Plotted are the individual replicates from the same simulations as in (a) and (b), respectively. Data are means ± SEM, n = 3 per condition, with SEMs and *p* values determined as in **Fig. 1g** in the main text.

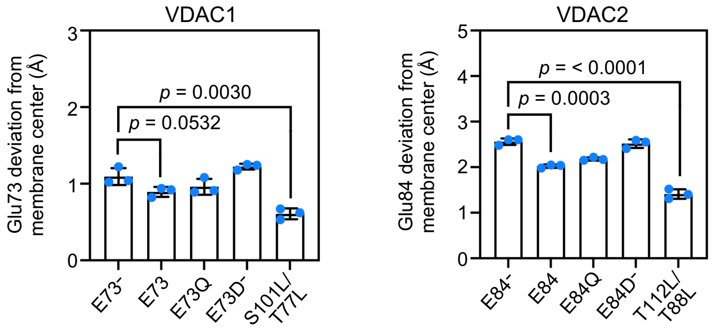


**Supplementary Figure 4. Exposure of the membrane-buried Glu to the cytosol is controlled by polarity and charge in the leaflet thinning vicinal region.**

Glu73/84 position relative to the center of geometry of the membrane bilayer for VDAC1/2 WT and mutants in an OMM-mimicking bilayer. Deviations from the membrane center indicate a higher exposure towards the cytosolic leaflet. The S101L/T77L and T112L/T88L variants were simulated with a deprotonated Glu73/84, respectively. Data are means ± SD, n = 3 per condition. *p* values were calculated by unpaired two-tailed *t* test.

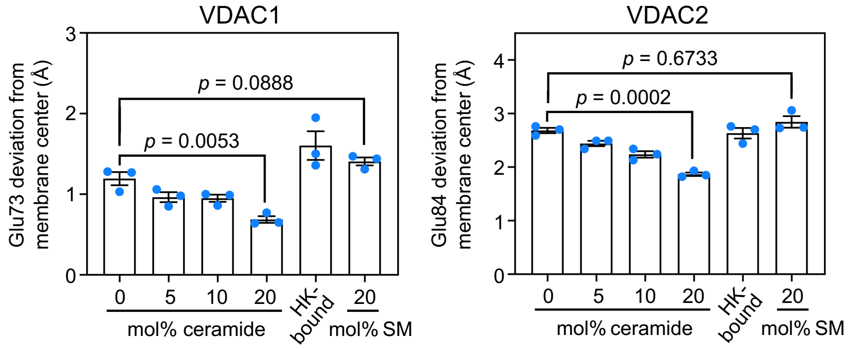


**Supplementary Figure 5. HKI and ceramides modulate the position of the membrane-buried Glu with antagonistic effects.**

Glu73/84 position relative to the center of geometry of the membrane bilayer for VDAC1/2 in an OMM-mimicking bilayer containing the indicate amount of $C_{16:0}$ ceramide or $C_{16:0}$ sphingomyelin. For the HK-bound state without ceramide, the same simulations that yielded **Fig. 1c,e** in the main text were used. Here, all frames in which HKI-Met1 was in direct contact with Glu73/84, respectively, were extracted and the position of the membrane buried Glu relative to the membrane center measured. Data are means ± SD, n = 3 per condition. *p* values were calculated by unpaired two-tailed *t* test.

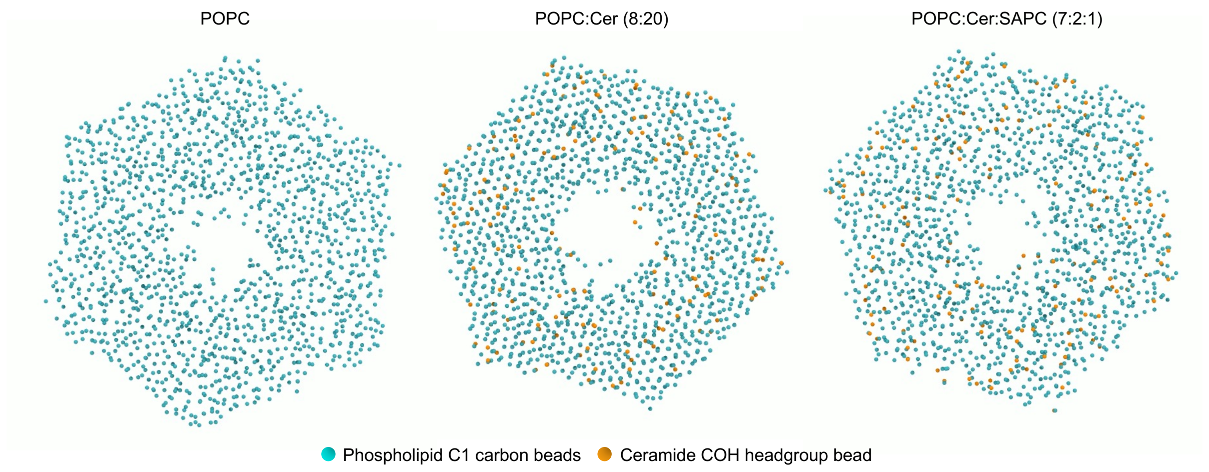


**Supplementary Movie 1. Ceramides cause the formation of a lipid ordered phase**

Side by side comparison of different membrane compositions and their respective fluidity. Shown are the phospholipid C1 carbon beads (*cyan*) and the ceramide COH headgroup beads (orange). VDAC channels are not shown for visual clarity. VDAC causes a distortion of the ordered membrane phase in the POPC:$C_{16:0}$ ceramide (80:20 mol%) membrane condition (*middle*) due to the inherent mismatch of its hydrophobic surface area (~2.4nm) with the membrane bilayer (~4nm). Length of the shown excerpts is approx. 150 ns for each condition.
